# In vitro reconstitution defines the mechanistic basis of HSET motor activity regulation by IntraFlagellar Transport proteins

**DOI:** 10.1101/2025.01.13.632783

**Authors:** Audrey Guesdon, Valérie Simon, Ron Siaden Ortega, Juliette Van Dijk, Julien Marcoux, Bénédicte Delaval, Benjamin Vitre

## Abstract

HSET is a mitotic kinesin essential for centrosome clustering in cells harboring supernumerary centrosomes. Work *in cellulo* revealed that IntraFlagellar Transport proteins (IFT) interact with the kinesin HSET to promote efficient extra centrosome clustering and subsequent cancer cell proliferation. However, whether and how IFT proteins regulate HSET activity is unknown. Using a reconstituted *in vitro* system combining purified HSET and IFT proteins with TIRF microscopy approaches, we identified a minimal subcomplex made of IFT52/IFT70 directly binding to HSET. We show that this binding induces HSET oligomerization promoting the formation of processive HSET complexes. We also show that HSET’s increased processivity upon IFT52/70 binding accounts for an increased ability to slide microtubules and to organize dynamic microtubule networks *in vitro*. Overall, this work shows that IFT proteins can directly promote the processive motility of a mitotic kinesin and provides a mechanistic explanation for the contribution of IFT proteins to efficient centrosome clustering.

## Introduction

Mitotic spindle bipolarity is essential to ensure accurate segregation of the genetic material in the two daughter cells during cell division. In animal cells, the formation of a bipolar mitotic spindle requires the establishment of polarized radial arrays of microtubules focused around the spindle poles. Those arrays, or asters, are subsequently separated and stabilized in a bipolar structure through the contribution of mitotic motors and microtubule-associated proteins ^1^.

Microtubule focusing at the spindle poles depends on the microtubule minus-end directed motor dynein in association with NuMA ^2–4^ and on microtubule minus-end directed motors of the kinesin-14 family including Ncd in drosophila ^5^ and HSET, also known as KIFC1, in human ^6^. Kinesin-14 motors are dimeric as demonstrated by structural and photobleaching experiments ^7,8^. They present conserved domain organization with an ATP-independent microtubule binding domain at their N-terminus, a coiled-coil domain in their central part and an ATP-dependent microtubule binding and motor domain at their C-terminus ^9^. Kinesin-14 are non-processive motors but can become processive through oligomerization ^8,10,11^. *In vitro*, kinesin-14 motors display a minus-end directed activity ^6,12,13^. Their two microtubule binding sites allow them to crosslink microtubules and focus their minus ends ^14,15^ favoring aster formation ^16^. In cells, the contribution of kinesins-14 to spindle pole formation and maintenance varies between organisms or cell type. For instance, Ncd is essential for aster formation and spindle pole focusing in drosophila mitotic cells ^17^ while HSET has only mild effects on regulating spindle size in human somatic cell ^18^. However, HSET is important for the assembly of acentrosomal mouse oocyte mitotic spindle ^6^ and is essential in cells harboring multiple centrosomes. Indeed, in cancer cells that frequently harbor supernumerary centrosomes ^19^ HSET is required for extra centrosome clustering leading to pseudo-bipolar mitotic spindle organization ^20^. Upon HSET depletion, cancer cells harboring supernumerary centrosomes fail to cluster their centrosomes leading to catastrophic mitotic errors and subsequent cell death ^20,21^.

Diverse mechanisms fine-tuning the activity of kinesin-14 motor activity throughout mitosis were identified over the years. First, HSET abundance is regulated through Cdk1 dependent phosphorylation within a D-box at its N-terminus which prevents its degradation by APC/C^CDH1^ during early mitosis ^22^. Second, HSET motor activity *per se* can be positively or negatively regulated. For example, Xenopus kinesin-14 XCTK2 activity is negatively regulated through its interaction with importins α/β ^23,24^. *In vitro*, individual HSET motility can be promoted by low ionic strength ^25^ or by applying a traction force on them ^26^. Finally, HSET clustering or oligomerization either through its N-terminal tail binding to soluble tubulin ^8^ or through binding to a bead ^27^ promotes its motility. Similarly, HSET oligomerization through its association with the plus-end tracking protein Mal3 could promote its activity ^28^.

Previous work carried out in cellular systems also identified IntraFlagellar Transport Proteins (IFT) as a novel class of proteins interacting and functionally regulating HSET to allow efficient clustering of extra centrosome ^29^. Indeed, this work showed that HSET can directly interact with a core complex made of IFT46/52/70/88 proteins and that disrupting this complex by depletion of IFT52 or IFT88 results in extra-centrosome clustering failure in cells. IFT proteins are primarily described to form large macromolecular complexes called IFT trains, that function as cargo adaptors of ciliary proteins ^30^. In cilia, IFTs are organized in large molecular complexes composed of multiple unique IFT proteins. Those complexes are assembled repeatedly to form ciliary trains ^31,32^. A core complex made of IFT46/52/70/88 is essential for the stability of the whole IFT complex and for cilia formation ^33^. Interestingly, *in vitro* work also identified regulatory roles for a complex formed of the *C. elegans* orthologs of IFT46/52/70/88 proteins on the ciliary kinesin OSM-3 ^34^. This study showed that *C. elegans* IFT70 ortholog is essential for the interaction with OSM3 but the mechanism by which IFT proteins regulate the activity of OSM 3 has not been elucidated. Similarly, the mechanism by which IFT46/52/70/88 can regulated the mitotic kinesin HSET are not known.

Using *in vitro* reconstituted assays, we identify here a minimal IFT complex composed of IFT52/70 that directly interacts with the human kinesin HSET. This interaction promotes the processive activity of the motor and stimulates its microtubule sliding activity. We also show that HSET’s increased processivity upon IFT52/70 binding enhances HSET bundling and clustering capacity. This mechanistically explains the role of IFT proteins/HSET interaction in the extra-centrosome clustering observed *in cellulo* ^29^.

## Results

### HSET binding to a minimal IFT52/70 complex stimulates its processive activity

We previously showed that IFT proteins are functionally required for efficient extra centrosome clustering by the kinesin HSET in cells with supernumerary centrosomes ^29^. We also showed that HSET interacts *in vitro* with a purified recombinant IFT-B core subcomplex made of IFT46/52/70/88 from *Chlamydomonas reinhardtii* ^29^. Taking advantage of SNAP-tagged mammalian IFT protein purification from baculovirus-infected insect cells and in vitro complex reconstitution, we sought to characterize more precisely whether and how IFT proteins modulate HSET activity. Using HSET GFP-trap pull-down, we first showed that a minimal IFT complex made of SNAP-tagged IFT52 and IFT70 directly interacts with HSET (Fig. 1a, b, c). To confirm this interaction and assess its impact on motor activity, we then combined *in vitro* complex assembly and TIRF microscopy. GMPCPP-stabilized microtubules were attached to a coverslip in PLL-PEG passivated imaging chambers. HSET with or without IFT52/70 were flown-in to allow the observation of HSET or IFT-HSET complexes. When HSET was incubated with IFT52/70 we observed processive particles containing both HSET and IFT52/70 (labelled with Alexa fluor 647 on their SNAP tag). This confirms that a subcomplex of IFT proteins made of IFT52/70 interacts with HSET when the kinesin was bound to microtubules (Fig. 1d, e, Supplementary Fig. 1a and Movie 1). As expected HSET alone mostly displayed a diffusive behavior with rapid bidirectional movement and frequent directional switch (Fig. 1d, Supplementary Fig. 1a and Movie 2). However, IFT52/70 binding to HSET significantly increased processive HSET runs with an average of 0.046± 0.001 processive event per µm of microtubules per minute compared to 0.010± 0.002 in HSET alone condition (Fig. 1e). As a negative control, HSET was incubated with IFT46, which displayed a weaker binding to HSET compared to IFT52/70 in pull-down assay (Fig. 1a, b, c) and did not significantly increase the rate of processive events (Fig. 1e, Movie 3). Of note, neither IFT46 alone nor IFT52/70 interact with microtubules in absence of HSET (Supplementary Figure 1b). In addition to increase the occurrence of processive events, IFT52/70 binding to HSET also significantly increase the processivity of HSET particles with a 25 % increase in run length (3.79± 0.23 µm versus 4.73± 0.15 µm, Fig. 1f). Altogether, these results show that a minimal IFT subcomplex made of IFT52/70 directly binds to HSET and specifically promotes its processive motility.

**Figure 1.**
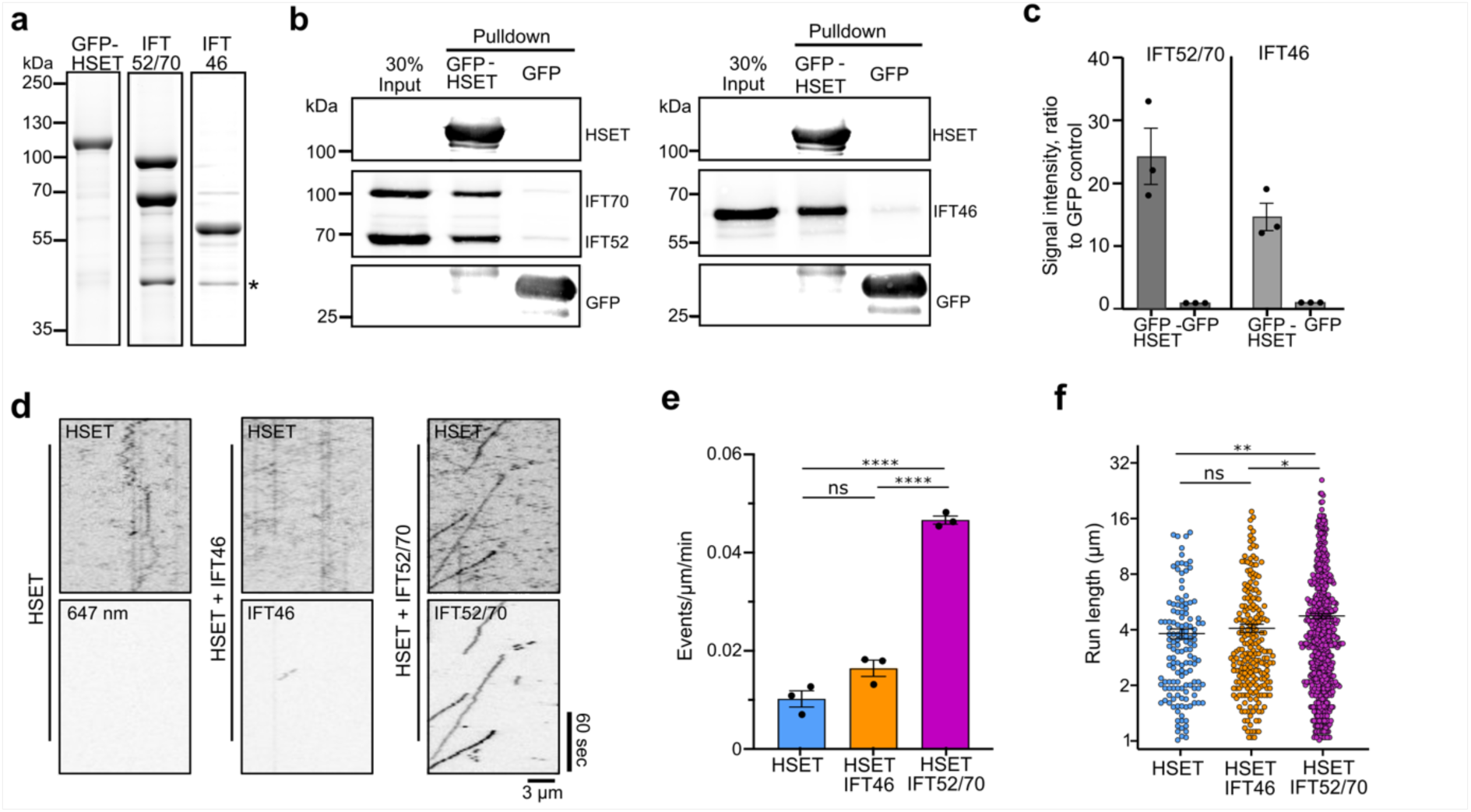
HSET binding to a minimal IFT52/70 complex stimulates its processive activity. **a.** Coomassie blue staining of purified GFP-HSET, SNAP-IFT52/SNAP-IFT70 and SNAP-IFT46. Asterisk indicates the HRV-3C protease used to cleave IFT proteins from their Strep-Tags during purification. **b.** Western-blot of a GFP-Trap pull down of IFT52/70 dimer using GFP or GFP-HSET (left) and GFP-Trap pull down of IFT46 using GFP or GFP-HSET. **c.** Quantification of the ratio of IFT52/70, IFT46 proteins pulled-down by GFP-HSET to a GFP alone control. Bars represent the average of three independent experiments error bars s.e.m. **d.** Kymographs representing individual GFP-HSET (0.5 nM) particles moving along GMPCPP stabilized microtubules in absence or presence of IFT proteins (IFT46 or IFT52/70 at 1nM). **e**. Graph representing the number of processive events of GFP-HSET particles alone or associated to the indicated IFT proteins. Each experimental replicate is a value obtained by dividing the total number of events observed in one condition by the total length of microtubule on which the events occurs divided by the imaging time (unit: events/µm/min). Three independent experiments were analyzed and are represented by black dots on the graph. **f.** Graph showing the processivity of HSET particles alone or associated to the indicated IFT proteins. For e. and f., three independent experiments were analyzed. A total of 137 events were quantified in control, 223 in HSET-IFT46 and 630 in HSET-IFT52/70. Bars indicate means and error bars S.E.M. * P<0,05; **p<0,01; ****P < 0.0001; unpaired t-test.

### IFT52/70 binding stimulates HSET processive motility through its oligomerization

To test if IFT binding to HSET follows a cooperative binding model, a range of IFT52/70 concentrations was incubated with a fixed concentration of HSET (0,18 nM). Increasing IFT52/70 to HSET ratio, increasingly promoted HSET processive motility (Fig. 2a) indicating that HSET’s increased processivity upon IFT52/70 binding follows a cooperative binding model where the motor is activated by the binding of multiple partners ^35,36^. To get further insights into the nature of HSET-IFT52/70 complexes we then used mass photometry ^37^. We analyzed IFT52/70 and observed two major peaks (Fig. 2c). The first one at 67±13 kDa could correspond to background signal from the buffer (Fig. 2b), contaminants such as HRV-3C present in IFT purification (* Fig. 1a), monomeric SNAP-IFT52 (expected molecular weight (MW) of 67 kDa) or SNAP-IFT70 (expected MW of 95 kDa) and their degradation peptides. The second peak at 148±30 kDa was consistent with IFT52/70 assembled as a dimer with an expected (MW) of 162 kDa (Fig. 2c). We next analyzed GFP-HSET (Fig. 2d), and observed two major peaks. The first one at 66±15 kDa corresponded to buffer background signal and contaminants (degradation peptides of GFP-HSET). The second peak, with a measured MW of 232±34 kDa was consistent with a dimer of GFP-HSET with an expected MW of 210 kDa. Finally, to confirm that they can form a complex as suggested by the GFP-Trap pull-down (Fig.1b), we analyzed an incubation of IFT52/70 with GFP-HSET (Fig. 2e). The analysis revealed four peaks: a first one at 60±12 kDa, corresponding to background signals and contaminants (see description of Fig 2c and 2d), a second peak at 153±20 kDa corresponding to IFT52/70 dimer and a third peak at 265±29 kDa corresponding to GFP-HSET dimer. The fourth peak measured at 525±51 kDa was consistent with a complex made of one dimer of GFP-HSET and two dimers of IFT52/70 with and expected MW of 534 kDa based on the amino acid sequence (or calculated at 571±29 kDa if inferred from the values of peak two and three). This result was observed in three independent replicates. Altogether, this mass photometry analysis shows that both IFT52/70 and GFP-HSET are present as dimers in solution and that they can form a complex containing two IFT52/70 per GFP-HSET in vitro (Fig. 2f). We next wondered if HSET bound to multiple IFT52/70 dimer could allow for the formation of complexes containing multiple IFT52/70 and HSET dimers (Fig. 2f). While some additional counts were measured by mass photometry around 760 kDa potentially indicating higher order oligomers, they were not identified as a proper peak due to the small number of counts and poor gaussian fit (Supplementary Fig. 1c). We thus analyzed the number of HSET motors present in the processive particles identified in TIRF microscopy (Fig. 1d). Since HSET oligomerization was previously shown to increase its capacity to do processive runs ^8,27^, we expected to found an increased number of HSET motors in the processive particles. To test this hypothesis, we measured the fluorescence intensity of processive HSET particles when HSET is alone or in the presence of IFT52/70. Fluorescence signal intensity measurements of a dimeric GFP-HSET (see methods and Supplementary Fig. 1d) were used to quantify the number of GFP molecules per active HSET particle. At the population level, IFT52/70 binding to HSET increased active particle median signal by 66 % (fluorescence intensity of 1349 a.u vs 2246; Supplementary Fig. 1e) suggesting that it favors HSET oligomerization on microtubules. To visualize this effect at the level of individual particles, the distribution of particles according to the number of HSET dimer per particle was analyzed in a frequency distribution histogram (Fig. 2g). In control condition, the majority of processive particles contained one or two HSET dimers (88 particles corresponding to 80 % of all processive particles) with the largest group containing a single HSET dimer (49 particles corresponding to 39,5 % of all processive particles). Strikingly, in presence of IFT52/70, the majority of HSET processive particles on microtubules contained more than two HSET dimers (336 particles corresponding to 59,4 % of all processive particles) indicating increased oligomerization in this condition. To confirm the increased oligomerization of HSET in presence of IFT52/70 a statistical analysis of the data was performed. Fit curves using non-linear regression with a gaussian fit were calculated and multiple unpaired t-test analysis on the best fit values for the mean number of GFP-HSET dimer per particle indicated a significant difference (p<0.05). Similarly, a Chi ^2^ test on the distribution of the data between HSET and HSET-IFT52/70 conditions showed a significant difference (p<0.0001). Altogether this confirms that IFT52/70 binding to HSET significantly increases HSET oligomerization. These results suggest that IFT52/70 binding to HSET promotes its processive motility by favoring its oligomerization. This is consistent with a previous study describing an increased HSET processivity through its oligomerization by soluble free tubulin ^8^.

**Figure 2.**
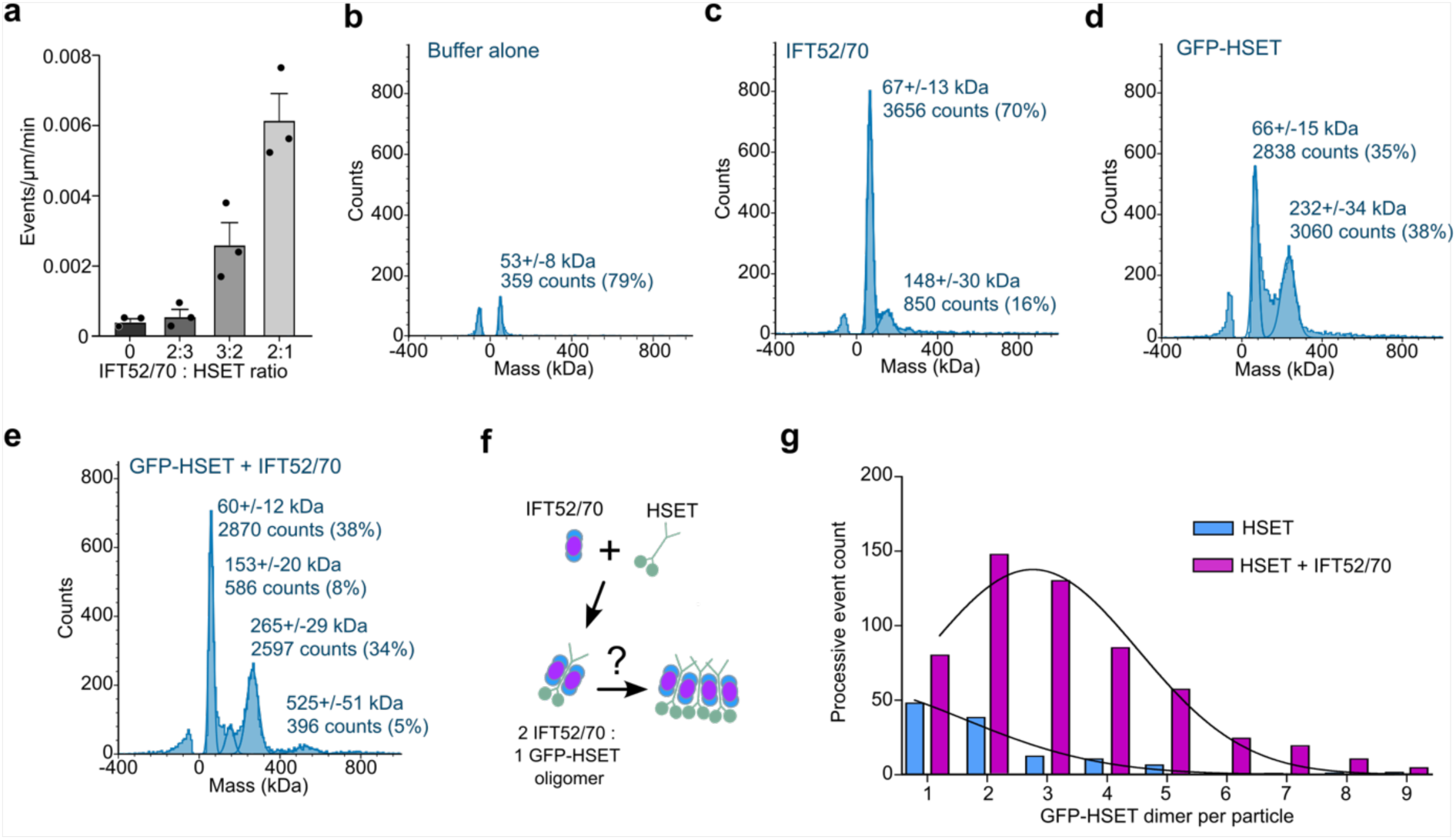
IFT52/70 binding stimulates HSET processive motility through its oligomerization. **a.** Graph representing the number of processive events of GFP-HSET particles alone or associated to IFT52/70. GFP-HSET concentration is fixed at 0.18 nM. IFT52/70 concentration is 0.12; 0.27 and 0.36 nM for 2:3, 3:2 and 2:1 ratio respectively. Three independent experiments were analyzed. Bars indicate means and error bars S.E.M. **b.** Mass photometry analysis of the buffer used in the experiment showing background signal. **c.** Mass photometry analysis of IFT52/70 (5 nM). **d.** Mass photometry analysis of GFP-HSET (2.5 nM). **e.** Mass photometry analysis of IFT52/70 and GFP-HSET at a 2:1 molar ratio (5 nM IFT52/70, 2.5 nM GFP-HSET). For all mass photometry data, text and numbers on the figures indicate the measured molecular weight and the number of counts of a specific peak. Percentage represent the proportion of total counts. **f.** Schematic showing a potential complex conformation made of IFT52/70 and GFP-HSET. **g**. Frequency distribution histogram indicating the number of processive particles depending on the number of GFP-HSET dimer per particle (dataset corresponding to Figure 1e). 124 events displayed in control condition and 566 in HSET-IFT52/70 condition. Data with more than 9 dimers are not plotted, see Supplementary Figure 1e for the full range of particle distribution. Black curves represent non-linear regression using a gaussian fit for HSET and HSET-IFT52/70 datasets.

### IFT52/70 binding to HSET favors its directional activity and its accumulation at microtubules minus ends

Analysis of processive HSET particles revealed that they frequently remain attached at microtubule ends in the presence of IFT52/70 (Fig. 3a). This suggested that IFT52/70 binding to HSET may favor HSET processivity and directional movement toward microtubule minus-ends, as expected from kinesin 14 directionality ^6,8,12,14^, subsequently leading to its accumulation. To validate the directionality of HSET runs upon IFT52/70 binding, we compared HSET localization to a plus-end directed kinesin Kip2-RFP known to accumulate at microtubule plus ends ^38^. As expected when successively injecting Kip2 and HSET with ITF52/70 in the imaging chamber, both HSET and IFT52/70 accumulated at the opposite end of microtubules compared to Kip2-RFP (Fig. 3b). This confirms HSET-IFT52/70 minus-end accumulation. To quantify HSET minus-end accumulation, we incubated 10 times more HSET (5 nM) with or without IFT52/70 (10 nM) and measured fluorescence intensity along microtubule length. At this higher concentration, HSET clearly accumulated at microtubule minus-ends in the presence of IFT52/70, forming minus-end comet-like structures referred to as “minus-end accumulation” later on (Fig. 3c, d). While a small accumulation of HSET was detected with the motor alone due to its occasional processivity (Fig. 1e), a 1.7-fold increase of HSET minus-end accumulation was observed upon incubation with IFT52/70 (1.38 versus 2.34 fluorescence ratio unit to control lattice, Fig. 3e). This showed that IFT52/70 binding to HSET promotes HSET particles processive motility toward microtubule minus-ends ultimately leading to their accumulation.

**Figure 3.**
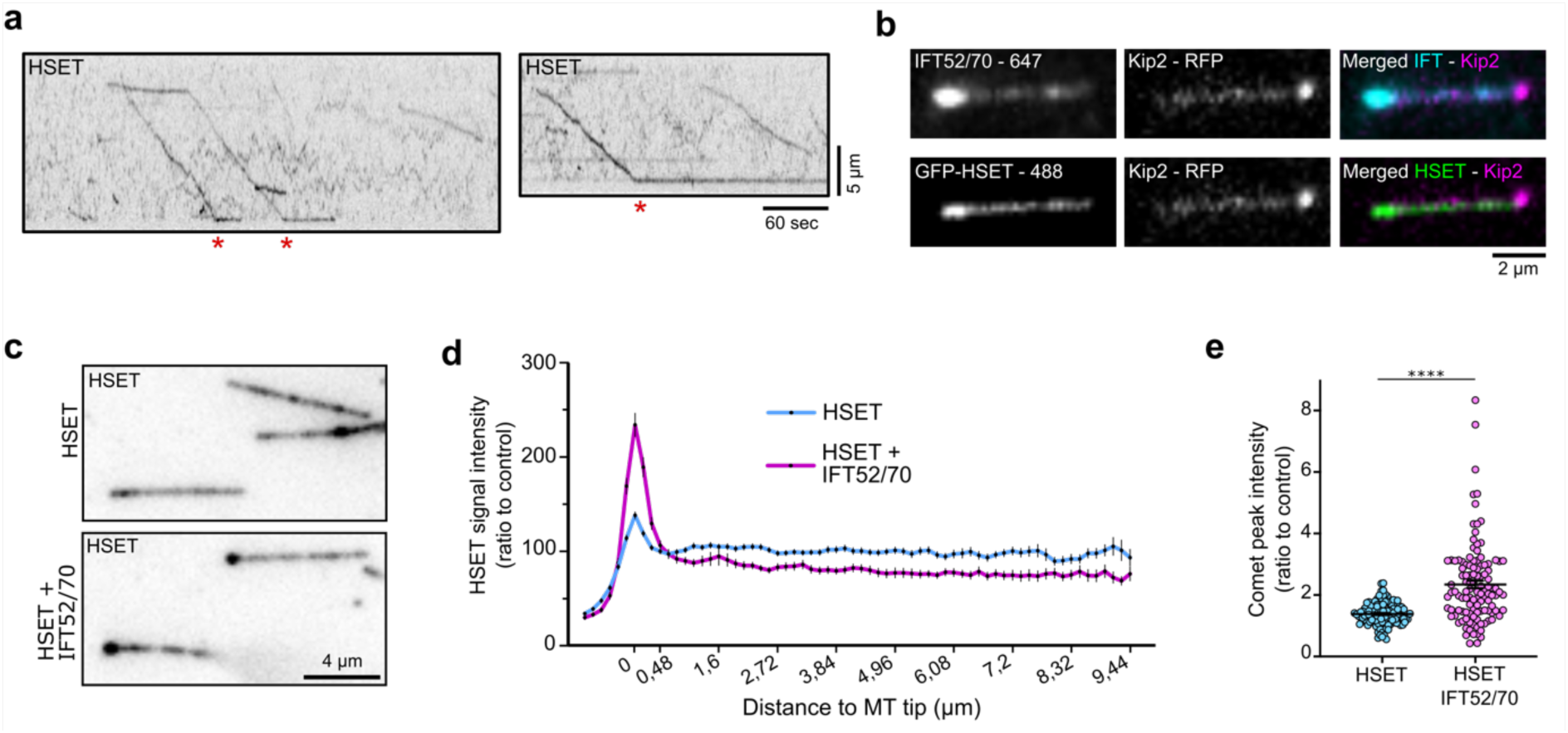
IFT52/70 binding to HSET favors its directional activity and its accumulation at microtubules minus ends. **a**. Representative images of GFP-HSET particles reaching a microtubule end and staying attached to it. HSET 0.5 nM and IFT52/70 1 nM. The red asterisks indicate the time when particles reach the end. **b**. Representative image of microtubules decorated with Kip2-RFP, and GFP-HSET colocalizing with IFT52/70. The two motors accumulate at opposite microtubule ends allowing the identification of plus and minus microtubule ends. **c**. Image of GFP-HSET decoration of microtubules after 2 minutes of incubation with GFP-HSET alone (5 nM) or GFP-HSET (5nM) plus IFT52/70 (10 nM). **d**. Graph representing GFP-HSET fluorescence signal intensity along microtubule length relative to the average signal of GFP-HSET along the lattice of microtubule in control condition. **e**. Graph representing GFP-HSET fluorescence signal intensity at microtubule minus-end relative to the average signal of GFP-HSET along the lattice of microtubule in control condition. Three independent experiments were analyzed. N=111 events in control condition and 115 in HSET-IFT52/70 condition. Bars indicate means and error bars S.E.M. *\*\*\*\*P < 0.0001*; Unpaired Student’s t-test.

### IFT52/70 binding to HSET increases its bundling and sliding activity

Functionally, kinesins-14 are required to remodel microtubule networks by crosslinking/bundling and sliding anti-parallel microtubules towards their minus ends both in cells ^18,39^ and *in vitro* ^14,15,40^. We thus tested whether HSET binding to IFT52/70 could affect its bundling and sliding activity. To visualize microtubule bundling and sliding upon collective HSET activity, biotinylated GMPCPP stabilized microtubules dimly labelled with Atto565 were attached to the coverslip of the imaging chamber (template microtubules, supplementary Fig. 2a). Bright non-biotinylated microtubule seeds were then added with IFT52/70 alone, with HSET alone or with HSET pre-incubated with IFT52/70 and monitored using TIRF microscopy. With IFT52/70 alone we did not observe any event of microtubule seeds bundling and sliding (Supplementary Fig. 2a). As expected, HSET alone induced free seeds bundling and sliding (Fig. 4a, Supplementary Fig. 2a, b and movies 4, 5). Of note, IFT52/70 colocalized with HSET between bundled seeds indicating that IFT52/70 direct binding to HSET is maintained when HSET crosslinks adjacent microtubules (Fig. 4a, b). In the presence of IFT52/70 (Fig. 4a, Supplementary Fig. 2b and movies 4, 5), the total number of bright microtubule seeds bundled with dim template microtubules was significantly increased showing that IFT52/70 binding to HSET increased its bundling capacity by 30 % (0.043± 0.003 events per µm of template in the control versus 0.058± 0.004 with IFT52/70; Fig. 4c). Among the bundled seeds, IFT52/70 binding to HSET also strongly increased the number of mobile seeds sliding on template microtubules (70% increased from 0.018± 0.001 events per µm of template in the control to 0.031± 0.002 in the condition with IFT52/70; Fig. 4d). The average speed of those moving seeds also increased in the presence of IFT52/70 with a distribution of individual seed speed shifted to higher values and an average seed speed increased by 40 % (0.458± 0.02 µm/min in the control versus 0.650± 0.02 µm/min with IFT52/70; Fig. 4e, f). Previous studies have shown that microtubule gliding or sliding speed can be differentially impacted depending on the concentration or density of HSET ^15,27^, XCTK2 ^16^ or Ncd ^41^ present in the assay. We thus assessed whether higher density of HSET motor in the presence of IFT proteins could account for this increased bundling and sliding capacity. To quantify HSET density on sliding microtubule seeds, we measured the ratio of GFP-HSET fluorescence signal to tubulin Atto565 signal (Fig. 4g) and showed that HSET accumulates at higher density on sliding microtubules in presence of IFT52/70 (ratio value of 0.43±0.13 in control versus 0.56±0.20 with IFTs). Altogether, these results show that IFT52/70’s direct binding to HSET increases its bundling and sliding capacity and correlates with a higher density of motors. These results are also consistent with the increased motor processivity observed at the single particle level (Fig. 1 and Fig. 2).

**Figure 4.**
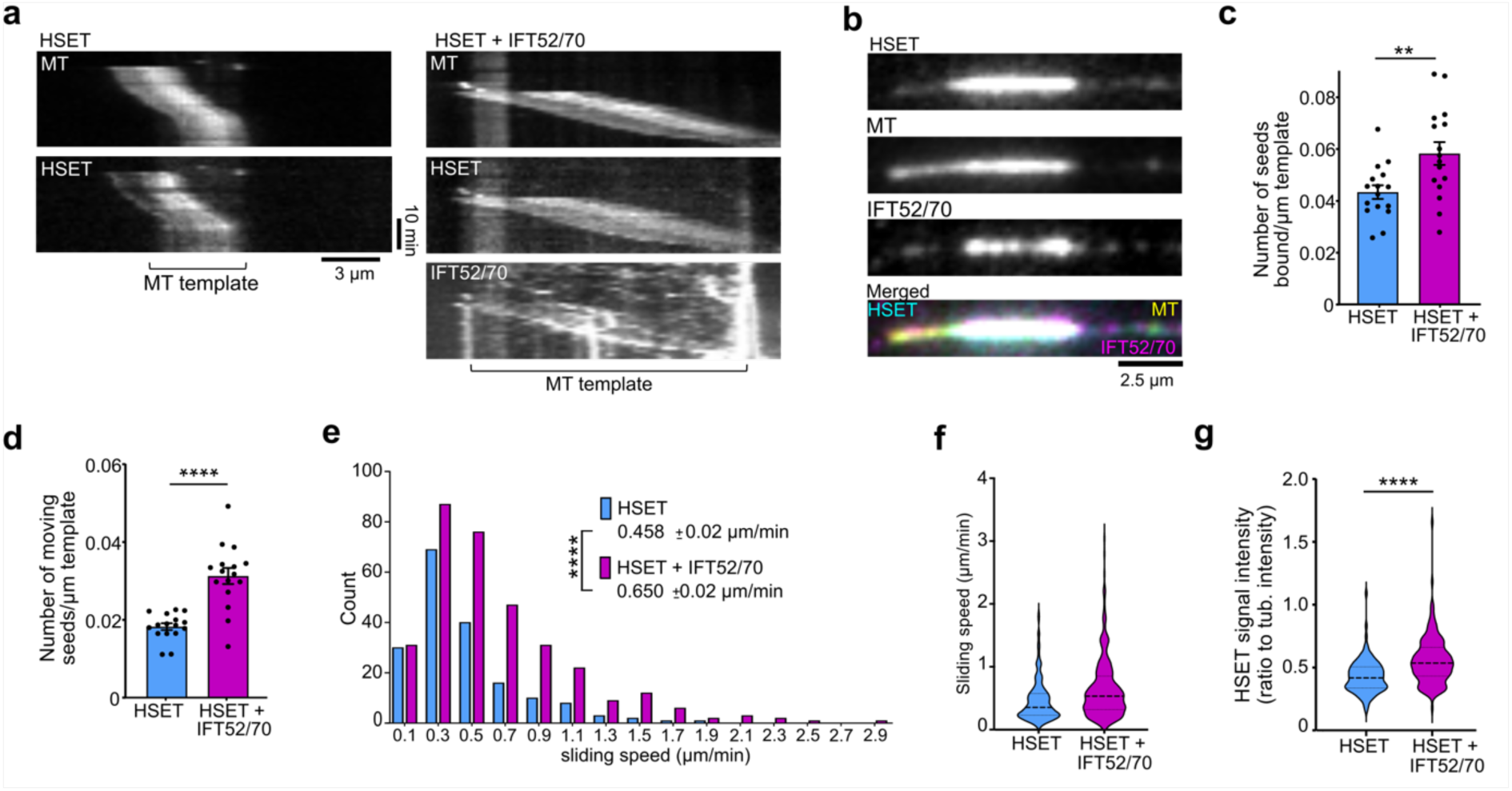
IFT52/70 binding stimulates HSET bundling and sliding activity. **a.** Kymographs of sliding of free microtubule seeds in the presence of HSET (50 nM) or HSET and IFT52/70 (100 nM). MT: microtubule. **b.** Representative image of HSET and IFT52/70 accumulation at the overlap between a free microtubule seed and the microtubule template in the presence of HSET and IFT52/70. The microtubule seed is the bright signal, the template is more clearly visible in the HSET channel which decorates microtubules evenly at 50 nM. **c**. Graph showing the total number of seeds bound per µm of template. **d**. Graph showing the number of moving seeds per µm of template. For **c.** and **d.** each point represents the measure from a single field of view. 16 fields of view from four independent experiments were analyzed for each condition. Bars indicate means and error bars S.E.M. **e**. Graph showing the speed distribution of individual microtubule seeds. Average speed was measured with N=180 for HSET and N=331 for HSET plus IFT52/70 from four independent experiments. **f**. Violin plot of the free microtubule seeds speed. All the events presented in panel d are represented. Bars indicate median and quartiles. **g.** Violin plot of the ratio of GFP-HSET mean fluorescent signal over tubulin mean fluorescent signal intensity on moving seeds. Bars indicate median and quartiles. N=116 for HSET and N=155 for HSET plus IFT52/70 from four independent experiments. \*\**P < 0.01* ; \*\*\*\**P < 0.0001* ; Unpaired Student’s t-test.

### IFT52/70 binding to HSET favors the formation of active microtubule networks

Minus-end accumulation of motor proteins such as dynein ^42,43^ or HSET ^44^ favors large scale microtubule network organizations into polarized aster structures. Moreover, it was shown experimentally and by simulation that crosslinking and sliding of microtubule minus-end-directed-motors such as HSET favors the formation of active microtubule networks that eventually evolve towards asters over time depending on motor and microtubule concentrations ^16,44–46^. We thus tested whether the increased accumulation of HSET at microtubule minus ends (Fig. 3) together with the increased sliding activity of HSET (Fig. 4) in the presence of IFT52/70 could increase HSET ability to organize active microtubule networks. Using an *in vitro* active microtubule network assay that can be monitored at low magnification (10 x objective), we showed that active microtubule network organization by HSET is increased in the presence of IFT52/70. Indeed, while microtubule bundles were already visible at the first imaging time point, one minute after mixing the components both in control and IFT52/70 conditions (Fig. 5a and movies 6, 7), they organized more rapidly into contractile microtubule networks (Fig. 5a, 10 min) and evolved faster towards radial microtubule organization or aster like structures in presence of IFT52/70 (Fig. 5a, 30 min). This can be precisely quantified by measuring overall image contrast over time which is a readout of microtubule structure compaction ^16^. In all conditions, contrast increased over time reflecting the microtubule network compaction, before tending towards a plateau at 30 minutes indicating a steady state in the network organization (Fig. 5b). However, increased compaction (i.e. more bundles) was already observed at the first timepoint (1 min after mixing the components) in the presence of IFT52/70 (1.017± 0.027 contrast a.u. in control versus 1.39± 0.057 with IFT52/70) and continuously increased until 30 minutes (2.68± 0.21 contrast a.u. in control versus 4.49± 0.27 with IFT52/70) reflecting a faster and stronger organization of the active microtubule network into aster-like structures in the presence of IFT52/70. We then measured the slopes of the linear parts of the curves (between 8 and 11 minutes) as a proxy of microtubule network compaction speed (Fig. 5c) and found that the presence of IFT52/70 increased 2.3 times microtubule compaction speed (slopes going from 0.07 to 0.16) confirming the faster contraction. We controlled that IFT52/70 alone did not organize active microtubule networks (Supplementary Fig. 3a), confirming the absence of bundling and sliding observed in TIRF sliding assay (Supplementary Fig. 2a). To further control for specificity and ensure that protein crowding alone was not responsible for the observed effect, we used IFT46, which does not increase HSET activity (Fig. 1). We also used SNAP tag as an additional negative control that does not bind to GFP-HSET in pull-down assay (Supplementary Fig. 3b, c, d). A minimal effect of IFT46 and SNAP was observed on compaction speed (Fig. 5c). This indicates that protein crowding may slightly influence compaction but confirms that IFT52/70 specifically favors microtubule network compaction. Importantly, HSET and IFT52/70 are present along the contractile microtubule network and accumulate at the center of the asters by the end of the imaging period (Fig. 5d) consistent with a role of their minus-end accumulation in active network organization and aster-like structure formation.

**Figure 5.**
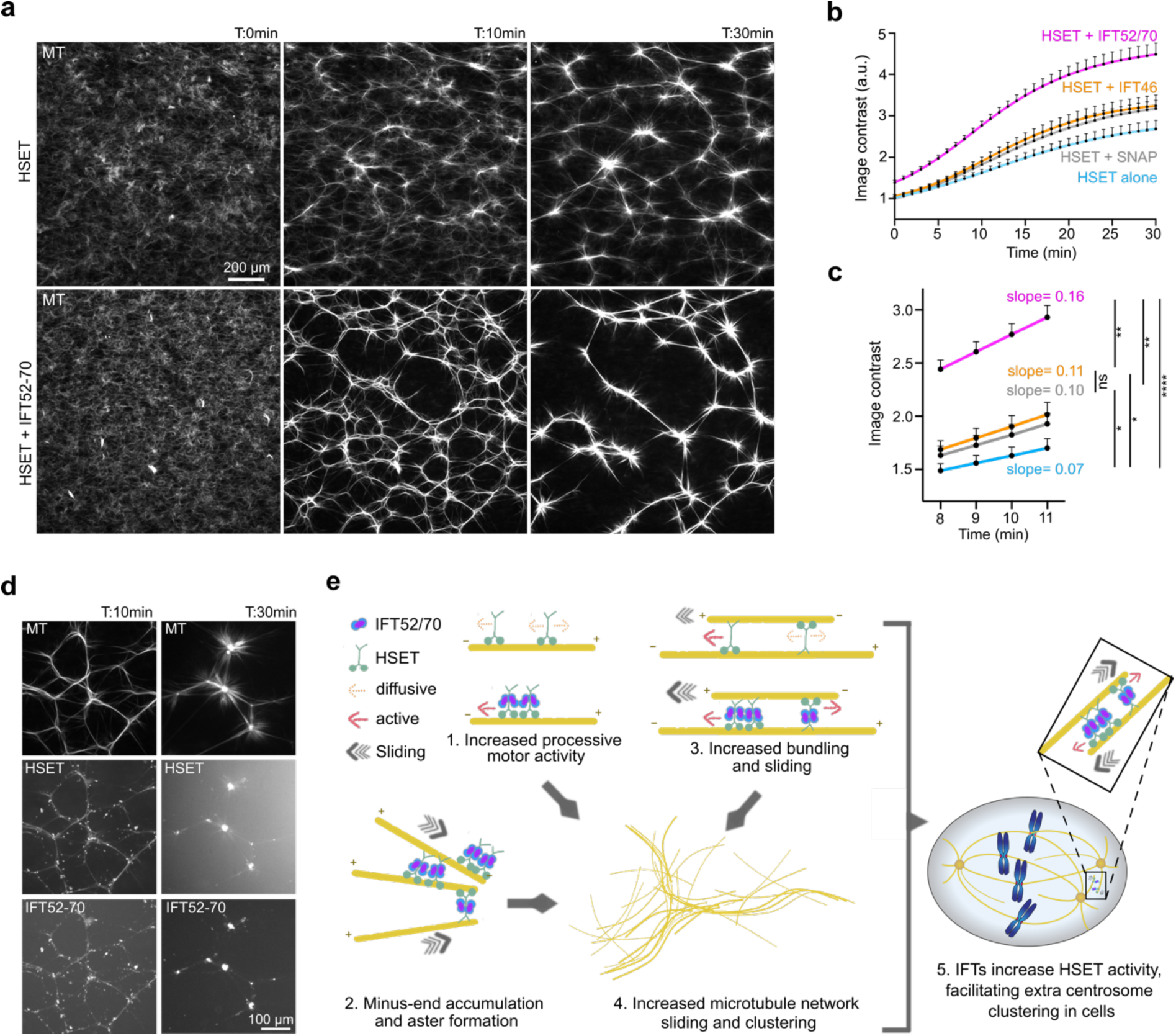
Increased HSET activity upon IFT52/70 binding favors the formation of active microtubule networks. **a.** Still images from live imaging of GFP-HSET (50 nM) forming contractile microtubule networks in absence or presence of IFT52/70 (100 nM) at 0, 10 and 30 min of incubation. MT: Microtubule. **b**. Graph showing image contrast measurements over time as a readout of microtubule compaction. **c**. Graph showing the values of image contrast (same values as panel b) between 8 and 11 minutes. The slopes of the curves were obtained using a simple linear regression. For panel b and c, points represent the average of a minimum of three independent experiments, errors bars S.E.M. N=27 individual measurements for HSET alone and HSET plus IFT52/70, N=26 for HSET plus IFT46 and N=14 for HSET plus SNAP. \**P < 0.05; **P < 0.01; ****P < 0.0001*; Unpaired Student’s t-test. **d**. Representative image of contractile fiber decoration in the tubulin, HSET and IFT52/70 channels in the HSET + IFT52/70 condition at 10 and 30 minutes. The same field of view is represented in the three channels. **e.** Model for IFT52/70 regulation of HSET activity as single particles or during microtubule network organization. IFT52/70 binding to HSET stimulates its processivity through oligomerization-(1) and its microtubule minus-end accumulation (2). It also increases HSET microtubule sliding activity (3) and HSET capacity to organize active microtubule networks into aster like structures (4). This activity of IFT52/70 on HSET is comparable to the activity of IFT proteins in cells which facilitates the clustering of extra centrosomes through HSET activity (5).

## Discussion

Using purified proteins and *in vitro* reconstituted assays, this study shows that IFT52/70 directly binds to HSET leading to its oligomerization into particles that show an increased processivity compared to individual HSET motors that are mostly non processive (Fig. 5e). This leads to an increased capacity of the motor to slide antiparallel microtubules and to the formation of active microtubule networks evolving toward the formation of asters (Fig. 5e). This work therefore provides mechanistic clues to understand the contribution of IFT proteins to efficient centrosome clustering in dividing cells harboring extra-centrosomes ^29^.

In light of previous work showing that kinesin-14 can organize microtubule aster *in vitro* ^16,44,47^, this work provides a novel regulatory mechanism for the kinesin through its interaction with a minimal complex of IFT proteins. It therefore complements studies showing that other minus-end directed motors such as dynein ^2,42,48^ or even plus-end directed motor such as the Kinesin-5 Kif11 ^44^ or oligomers of conventional Kinesin-1 ^49^ can organize polarized microtubule networks *in vitro* mimicking mitotic network organization.

While IFT proteins have been shown to regulate the *C.elegans* ciliary kinesin 2 OSM-3 ^34^ this work is the first demonstration of the regulation of a human mitotic kinesin by IFT proteins. Stimulation of HSET’s processive activity by oligomerization was previously observed via diverse mechanisms including oligomerization with tubulin forming clusters containing multiple tubulin dimers and HSET dimers ^8^. Similarly, binding of multiple HSET to a single quantum dot can also favor HSET processive activity ^50^. In solution, using mass photometry, we found HSET dimer bound to IFT52/70 with a one-to-two ratio but oligomers containing multiples HSET were not detected (Fig. 2). However, in TIRF microscopy we detected oligomers containing multiple HSET dimers. This difference could be due to the fact that oligomers display more processive activity (Fig. 1 and Fig. 2) and therefore are detected in a TIRF assay which aim at identifying processive particles. Alternatively, binding of HSET-IFT52/70 complexes to microtubules, may favor their oligomerization and/or complex stability in TIRF experiment compared to complexes analyzed by mass photometry in solution. This hypothesis is consistent with recent *in situ* Cryo-EM characterization of anterograde IFT train in *Chlamydomonas* cilia indicating that IFT52/70/88 could participate in lateral interaction between IFT repeats within the trains ^32^. Interestingly, the fact that HSET‘s processive motor activity is only stimulated when there is an excess of IFT52/70 to HSET (Fig. 2a) is consistent with the hypothesis of IFT52/70 oligomerizing multiple HSET and indicates that HSET increased processivity in the presence of IFT proteins follows a cooperative binding model. Comparable cooperative activation was described for the kinesin-1 since its negative regulation is relieved by binding to its partner JIP3 in a dose dependent manner ^36^. We did also quantify numerous events of single HSET dimer bound to IFT52/70 doing processive runs. We thus cannot exclude that part of the activating effects of IFT proteins on HSET can be driven by other mechanisms than oligomerization. Low ionic strength (5 µM Kac) has been proposed to increase HSET motor activity ^25^ but it is likely not the case here since our experiments were done at a conventional concentration of 50 mM KCl in absence or presence of IFT52/70. Another mechanism regulating kinesin activity is autoinhibition ^51^ but HSET does not seem to be inhibited by head to tail interaction since its N-terminal tail deletion is not sufficient to form a constitutively active motor ^8^ and HSET appears in an “open” conformation when observed by electron microscopy ^52^. Recent data indicate that kinesin-14 motor Ncd N-terminal tail interaction with NuMA’s fly ortholog Mud increased its affinity for microtubules ^53^. It is thus possible that similarly, IFT52/70 binding to HSET induces conformational changes that promote its processive motor activity. Future biochemical and structural biology work will be required to address this hypothesis. Our study also demonstrates that the increased HSET activity upon IFT52/70 binding results in HSET accumulation at microtubule minus ends (Fig. 3) as well as an increased ability to slide microtubules (Fig. 4). Those two mechanisms have been described to drive different organization of active microtubule networks with end-binding favoring aster formation while sliding favors contractile bundle formation ^16,44,46,49^. Indeed, when we assessed the effect of IFT52/70 bound HSET on microtubule network organization we observed first the formation of contractile networks that evolved towards asters over time (Fig. 5). This transition is driven by the equilibrium between the concentration of microtubules and the density of motors present in the experimental chamber as demonstrated using simulation ^45,46^ and HSET increased sliding activity and microtubule end-on accumulation upon IFT52/70 binding accelerate the process.

Of note, it was previously shown that increased density of kinesins-14 along sliding or gliding microtubules tend to slow down the moving seed speed potentially due to interaction of the kinesin N-terminal tails with microtubules that acts as a brake^15,16,27^. This effect seems contradictory to our observations, but it was shown in a range of concentrations below 25 nM and tends to be reversed above 50 nM ^16^ which is consistent with our observations.

Interestingly, this increased activity of HSET bound to IFT52/70 leading to the coalescence of multiple microtubule seeds into polarized structures is reminiscent of the regulatory role of IFT proteins in HSET dependent clustering of extra centrosomes we described previously in cells ^29^. The *in vitro* set up used here thus provides mechanistic clues to explain, using a bottom-up approach, the phenotype previously described *in cellulo* regarding the regulatory activity of IFT proteins on HSET motor activity (Fig. 5e).

Since extra centrosomes are frequently found in cancer while they are extremely rare in normal cycling cells, targeting HSET activity has appeared as a good strategy to specifically kill cancer cells ^54^. Specific inhibitor of HSET ATPase activity were thus developed showing efficacy *in cellulo* ^55–57^. However, inhibitors of kinesins that target their ATPase activity often have off target activity, and resistance due to the appearance of point mutations is a substantial drawback in the development of such inhibitors ^58^. Targeting regulatory proteins that interact with kinesins to inhibit specific functions appears as an attractive therapeutic alternative and small molecule perturbing IFT proteins-HSET interaction could be a novel way to modulate HSET activity.

## Material and methods

### Protein expression and purification

GFP-HSET was expressed in Sf9 insect cells and purified as described in ^15^. Briefly, full-length N-terminal hexa-histidine-tagged GFP-HSET cloned in pOET1C-modified vectors (gift of S. Diez, TU Dresden) was expressed in Sf9 insect cells by infecting them with the appropriate baculovirus for 48h. Harvested cells were resuspended in lysis buffer (20 mM Hepes pH 7.2, 300 mM NaCl, 2 mM MgCl_2_, 10 mM β-mercaptoethanol, 10 % glycerol, 0.5 % Triton X100, 50 µg/ml DNAse I, 1 mM PMSF, EDTA-free protease inhibitors (cOmplete Roche), 20 mM imidazole and 0.1 mM Mg-ATP). Cells were lysed using dounce homogenization and crude lysates were centrifuged at 75 000g for 45 min at 4 °C (25 000 rpm, Type 50.2 Ti rotor Beckman), loaded on Ni-NTA resin (Qiagen, 750 µl for 100 ml of cells) and incubated for 1h at 4 °C. The resin was washed for 2h with HSET purification buffer (20 mM Hepes pH 7.2, 300 mM NaCl, 2 mM MgCl_2_, 5 mM β-mercaptoethanol, 0.1 mM Mg-ATP, 10 % glycerol) containing 30 mM imidazole. Proteins were eluted in HSET purification buffer containing 200 mM imidazole (70 µl elution buffer for 100 µl resin). For IFT proteins, cDNA of human IFT46, mouse IFT52 and human ITF70 were cloned in 438-SNAP-V1 (addgene 55222) vector using LIC cloning as in ^59^. IFT52 and IFT70 were cloned in the same vector with their own polyhedrin promoter so they could be co-expressed together. The functionality of a mIFT52 within human IFT complex was previously validated in a rescue experiment in human cells ^29^. Both IFT52 and IFT70 have a twin strep tag and a SNAP tag in N-terminus. Baculoviruses were obtained using the pFastBac system. Proteins were expressed in Sf9 cells by infecting them with the appropriate baculovirus for 48h. Harvested cells were resuspended in lysis buffer (20 mM Hepes pH 7.2, 300 mM NaCl, 2 mM MgCl_2_, 10 mM β-mercaptoethanol, 10 % glycerol, 0.5 % Triton X100, 50 µg/ml DNAse I, 1 mM PMSF, EDTA-free protease inhibitors (cOmplete Roche) supplemented with 0.1 mg/ml avidin. Cells were lysed using dounce homogenization and crude lysates were centrifuged at 75 000g for 45 min at 4 °C (25 000 rpm, Type 50.2 Ti rotor Beckman). Clear lysates were loaded on Streptactin resin (100 µl for 100 ml of cells, StrepTactin Sepharose High Performance Cytiva) and incubated for 1h30 at 4 °C. The resin was washed with wash buffer (20 mM Hepes pH 7.2, 300 mM NaCl and 1 M NaCl, 2 mM MgCl_2_, 5 mM β-mercaptoethanol, 10 % glycerol) and proteins were eluted (strep tag cleavage by HRV-3C) and labelled simultaneously by incubating the resin O/N at 4 °C in elution buffer (20 mM Hepes pH 7.2, 300 mM NaCl, 2 mM MgCl_2_, 1 mM DTT, 10 % glycerol; 100 µl for 100 µl resin) supplemented with HRV 3C protease (Pierce, 30 units for 100 µl elution buffer) and fluorescent dye SNAP-Surface Alexa Fluor 647 (NEB). Eluates were finally loaded on a biotin and dye removal spin column (Zeba, Thermo). Proteins were snap frozen in liquid nitrogen and stored at −80 °C until use. Protein size, purity and concentration were checked and measured on SDS-PAGE gel. Tubulin was purified from porcine brain using the high molar Pipes buffer protocol ^60^. Purified tubulin was labelled with fluorescent dye or biotin using standard protocols from Mitchison’s Lab ^61^. Kip2-RFP was a gift from D. Portran expressed and purified as in ^38^.

### GFP trap Pull down assays

4 µg of purified GFP-HSET or GFP were incubated on 10 µl of GFP-trap beads for 1h at 4 °C. Beads were then incubated with IFT proteins (0.25 µM) in 20 mM Hepes pH 7.2, 150 mM NaCl, 2 mM MgCl_2_, 1 mM DTT, for 2h at RT (100 µl total reaction volume). Beads were washed 4 times and resuspended in Laemmli buffer for SDS-PAGE analysis. For western blots, GFP and GFP-HSET were revealed using a mouse monoclonal antibody against GFP (Proteintech, # 66002) while IFT46, IFT52, IFT70, SNAP were visualized using their labelling with SNAP-surface 647 (NEB).

### Mass Photometry

Microscope coverslips (24 x 50 mm, Epredia) were sequentially sonicated for 6 x 5 min in alternating baths of deionized water and isopropanol and then dried under a nitrogen stream. To perform a measurement, a reusable 6-well silicone gasket (Grace Bio-Labs) was placed onto a clean coverslip which was then placed into a TwoMP mass photometer (Refeyn). First a 15 μL droplet of reaction buffer was added to a well and used to adjust the focus. Then 5 μL of the protein solution was mixed into the droplet and 60 s movies were recorded using Refeyn AcquireMP v2024 R1.1. Acquired data was processed using Refeyn DiscoverMP v2024 R1, and a radiometry-molecular weight calibration was created using a NativeMark™ unstained protein standard with known MWs (ThermoFisher).

GFP-HSET and IFT52/70 were incubated at a 1:2 molar ratio (1.2 µM GFP-HSET and 2.4 µM IFT52/70) in reaction buffer (20 mM Na-Hepes pH 7.2, 150 mM NaCl, 2 mM MgCl2, 10 % glycerol, 5mM β-mercaptoethanol) at a final volume of 5 μL and incubated for 30 min at 30 °C in a water bath. The samples were then diluted to 10 nM of GFP-HSET and 20 nM of IFT52/70 with reaction buffer at 30 °C containing 150 mM NaCl. Mass photometry data was acquired within 30 s of the first dilution at a final concentration of 2.5 nM GFP-HSET and 5 nM IFT52/70 in the droplet. Experiments were replicated three time independently at 25 °C.

### TIRF microscopy chamber preparation

Experiments were performed in flow chambers assembled from a coverslip (Corning Cover glass Thickness ½ 24×50 mm; Cat No. 2980-245) attached to a glass slide (Knittel glass Starfrost 76×26 mm) with double-sided tape (NITTO cat EST-805, 50 µm thick). Flow chamber volumes were ranging from 5 to 8 µl depending on chamber width. Coverslips and glass slides were washed by sonicating in an isopropanol bath for 10 min, extensively rinsed with milliQ water (at least 10 “Hellendahl staining tank” volumes) and dried with argon. This step was followed by a plasma cleaning. Chambers were then passivated using PLL-PEG biotin (SuSos, 50 % labelled at 0.2 mg/ml final) for 5 minutes, washed with 2 to 3 volumes of BRB80 (1x), then flushed with neutravidin (0.25 mg/ml final) and incubated for 5 min. The chambers were finally washed with 2 to 3 volumes of BRB80 (1x) before injecting stabilized microtubules and the reaction mixture.

### Seed preparation for *in vitro* reconstituted assays

For HSET single particle dynamic assays and sliding assays, GMPCPP stabilized and biotinylated dimly labelled microtubules (i.e template microtubules containing 20 % biotin tubulin (0.6 µM) and 3 % Atto 565 tubulin (0.09 µM)) and non-biotinylated bright microtubules (i.e transported microtubules seeds containing 13 % of Atto 565 tubulin (2 µm)) were prepared as previously described ^62^. For active microtubule network assays, taxol-stabilized microtubules were prepared using a protocol adapted from the Mitchison lab to obtain a microtubule seed stock at a concentration of 25 µM tubulin.

### Single particle assay

HSET was incubated for 30 min at 30 °C with or without IFT proteins in incubation buffer (20 mM Na-Hepes pH 7.2, 150 mM NaCl, 2 mM MgCl2, 10 % glycerol, 5 mM β-mercaptoethanol, 100 µM Mg-ATP) at a concentration of 0.75 µM for HSET and 1.5 µM for IFT proteins. Template biotinylated microtubules in BRB80 were flushed into the chamber and incubated for 5 min in order to attach to the coverslip. Incubated proteins were flushed in the chamber in a reaction mix containing 20 mM NaPIPES pH 6.8, 1 mM EGTA, 7.5 mM MgCl_2_, 5 mM MgATP, 50 mM KCl, 1.5 mM GTP, 200 mM sucrose, 2 µM taxol, 0.25 mg/ml k-casein, 0.25 µM glucose oxidase, 0.064 µM catalase, 40 mM D-glucose, 17.5 mM β-mercaptoethanol. Final concentrations of proteins in single particle assay were 0.5 nM HSET, 1 nM IFT52/70 and in the minus-end accumulation assay, 5 nM for HSET and 10 nM for IFT52/70. Image acquisition was performed using an inverted Nikon wide-field microscope equipped with a ILAS2 TIRF module (Roper), a 100X Plan TIRF Apochromat 1.49NA oil objective, an EMCCD iXon 897 Ultra Andor camera (512*512, 16µm pixel size) detector, controlled by Metamorph (Multi-dimensional acquisition module). Images were acquired every 1.4 sec for 7.5 minutes in the 3 channels (488, 561, 642). For the minus end accumulation assay, after 2 minutes of incubation images were also acquired with 1.4 sec interval but only the first image was used for further quantification.

### Microtubule sliding assay

HSET was incubated for 30 min at 30 °C with or without IFT proteins in 150 mM NaCl Hepes buffer (incubation buffer) at a concentration of 0.75 µM for HSET and 1.5 µM for IFT proteins. Template biotinylated microtubules in BRB80 were flushed into the chamber and incubated for 5 min in order to attach to the coverslip. Incubated proteins were added to non-biotinylated microtubules in a reaction mix containing 20 mM NaPIPES pH 6.8, 1 mM EGTA, 7.5 mM MgCl_2_, 5 mM MgATP, 50 mM KCl, 1.5 mM GTP, 200 mM sucrose, 2 µM taxol, 0.25 mg/ml k-casein, 0.25 µM glucose oxidase, 0.064 µM catalase, 40 mM D-glucose, 17.5 mM β-mercaptoethanol, and injected in the chamber containing template microtubules. Final concentrations of proteins in the assay were 50 nM HSET, 100 nM IFT dimer, 120 nM biotinylated seeds, and 150 nM non-biotinylated seeds. Image acquisition was performed using an inverted Nikon wide-field microscope equipped with a ILAS2 TIRF module (Roper), a 100X Plan TIRF Apochromat 1.49NA oil objective, an EMCCD iXon 897 Ultra Andor camera (512*512, 16µm pixel size) detector, controlled by Metamorph (Multi-dimensional acquisition module). Images were acquired every minute for 30 minutes in the 3 channels (488, 561, 642). Three positions per condition were registered, all conditions were acquired in parallel chambers, at the same time.

### Microtubule active network formation assays

Experiments were performed in similar flow chambers than TIRF experiments (without coating for passivation). HSET (0.75 µM) was incubated with or without IFT52/70, IFT46 or SNAP (1.5 µM) at 30 °C for 30 min, in Hepes buffer (incubation buffer) containing a final concentration of 150 mM NaCl. The assay mix was prepared as following at room temperature: at the end of the incubation time, the appropriate volume of incubated sample was added to taxol -stabilized microtubule seeds, the sample was thoroughly mixed by pipetting up and down and quickly flushed into the chamber. Chambers were sealed with grease before acquisition. Images for all conditions (HSET alone, HSET/SNAP, and HSET/IFT protein conditions) were acquired at the same time in parallel chambers. The final concentrations used in the assay were 100 nM HSET, 200 nM IFT dimer, IFT46 and SNAP, and the equivalent of 2.5 µM tubulin of taxol stabilized seeds in an assay buffer containing 20 mM NaPIPES pH 6.8, 1 mM EGTA, 7.5 mM MgCl_2_, 5 mM MgATP, 50 mM KCl, 1.5 mM GTP, 200 mM sucrose, 2 µM taxol; 0.25 mg/ml k-casein, 0.25 µM glucose oxidase, 0.064 µM catalase, 40 mM D-glucose, 17.5 mM β-mercaptoethanol. Imaging was performed at 32 °C using an Inverted Nikon Ti-E wide-field microscope, a 10X Plan Apochromat 0.45NA air objective, and a sCMOS back-illuminated Prime95B Photometrics camera (1200*1200, 11 µm pixel size), controlled by NIS Elements (Nikon). Time-lapse wide-field fluorescence images were acquired at 1 min interval for a total duration of 30 min (31 frames). Three fields of view per chamber/condition were acquired on the appropriate channels.

### Minus-end directed movement determination

The study of the directionality of HSET-IFT protein complex was performed on fluorescent GMPCPP microtubules using TIRF microscopy. Microtubule seeds were first incubated into an observation chamber, in order to adhere to the coverslip. The plus end directed kinesin Kip2 was flushed into the chamber and incubated for 5 to 10 min at 30 °C in order to allow the formation of comets at microtubule plus ends. The chamber was then washed with BRB80 to remove excess of Kip2, before flushing preformed HSET-IFT52/70 complex diluted in microtubule self-organization assay buffer, at a final concentration of 50 and 100 nM, respectively. One image per second was acquired during 5 min to observe HSET-IFT microtubule minus-end accumulation (opposite end than Kip2 accumulation).

### Image Analysis

Image processing and analysis (cropping, rotating, brightness and contrast adjustment, color combining, and measurements) were performed using Fiji (ImageJ). For single particle analysis, kymograph of individual microtubules and velocity parameters were obtained using ImageJ Velocity Measurement Tool. Individual particle fluorescence (Supplementary Fig. 1e) is the average of the fluorescence signal of the first five time-intervals of each processive run. Values were corrected for background signal. To determine the number of GFP particles per track we first analyzed the average fluorescence decay of GFP-HSET particles attached to the imaging chamber that would bleach in one or two steps. The average intensity of particles that bleach in two steps was measured and set at 750 a.u. (Supplementary Fig. 1d). Individual fluorescence values of processive particles were divided by this value to determine the number of GFP-HSET dimer per moving particle (Fig. 2f). For sliding assays, the number of transport microtubules attached to and moving along template microtubules was measured manually. Sliding velocity was measured on kymographs, generated semi automatically with a homemade ImageJ macro including the plugin Kymograph Builder, and using the ImageJ Velocity measurement tool. Average fluorescence intensity of sliding microtubule was measured in the 488 and 561 channels using ROI overlapping the entire length of the moving seeds. The ratio of HSET signal to tubulin signal was calculated. This ratio was corrected with a correction factor corresponding to the ratio of HSET to tubulin on template microtubules without moving seeds, in the control condition, to take into account potential fluorescence signal variation in the two channels between different experiments. For active microtubule network assay image contrast was defined as the standard deviation of the fluorescent signal that was measured on the tubulin channel (561 nm). Normalization was done by dividing the original standard deviation values by the average control ratio (average of the 3 control fields at t = 0 Min) for each experimental replicate.

### Statistical analysis

The number of events per experiment used for statistical analysis is indicated in figure legends. Graphs were created and statistical analysis were done using GraphPad Prism software. P-values were calculated using unpaired two-tailed Student’s t-tests, multiple t-test or Chi^2^ test.

## Acknowledgments

We are grateful to D. Portran from the Liakopoulos team for providing reagents; to members of the Delaval and Liakopoulos team for fruitful discussions. This work was supported by the Agence National de la Recherche, grant number ANR-18-CE-0025-01 to BV and grant number ANR-19-CE13-0014 LUCELL to BD and by the Fondation pour la Recherche Médicale, “Equipe FRM EQU202103012567” to BD. We acknowledge the Biocampus Montpellier Facilities that contributed to the project: Montpellier Genomic Collection and Montpellier Ressources Imagerie (MRI). MRI, is member of the national infrastructure France-BioImaging supported by the French National Research Agency (ANR-10-INBS-04, «Investments for the future»).

## Data availability Statement

The source data underlying the figures are provided in Supplementary Data 1. All other data supporting the findings of this study are available within this article or can be obtained from the corresponding author upon request.

## Author contributions

Conceptualization: AG, BD and BV. Investigation: AG, VS, RSO, JVD, JM, BD, and BV. Methodology: AG, VS, RSO, JVD, JM, BD and BV. Supervision: BD and BV. Funding acquisition: BD and BV. Writing BD and BV with inputs from authors.

## Conflict of interest

The authors declare no conflict of interest

---

**Supplementary Figure 1.**
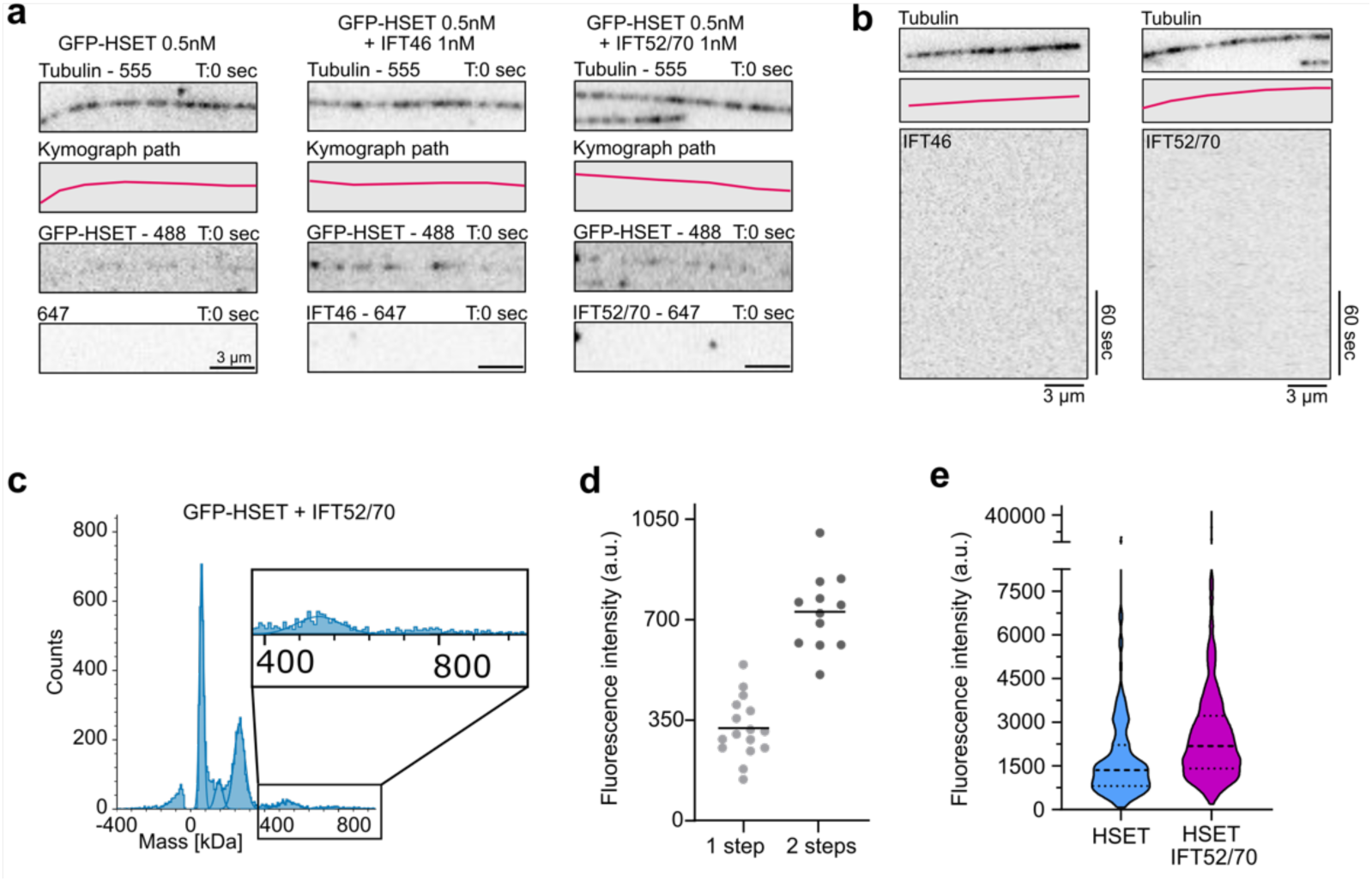
**a.** Images of the field of view from Movies 1, 2 and 3 used to generate the kymographs in Fig. 1d. **b.** Left: image of a field of view of a microtubule (top) use to generate a kymograph of IFT46 alone (bottom). IFT46 alone does not bind to microtubules. Right: image of a field of view of a microtubule (top) use to generate a kymograph of IFT52/70 alone (bottom). The IFT52/70 dimer alone does not bind to microtubules. **c.** Mass photometry analysis of GFP-HSET dimer (2.5 nM) incubated with IFT52/70 dimer (5 nM) as presented in Figure 2e with a zoom-in inset of the event between 400 and 960 kDa showing multiple counts between 720 and 800 kDa. **d.** Dot plot of the fluorescence signal of GFP-HSET particles bleaching in one or two steps allowing to identify the average fluorescence intensity of a GFP-HSET dimer (two GFP, two steps). Bars indicate average signal. **e**. Violin plot of GFP-HSET individual particles fluorescent signal. Bars indicate median and quartiles. N=127 for HSET condition and 591 for HSET-IFT52/70 condition.

**Supplementary Figure 2.**
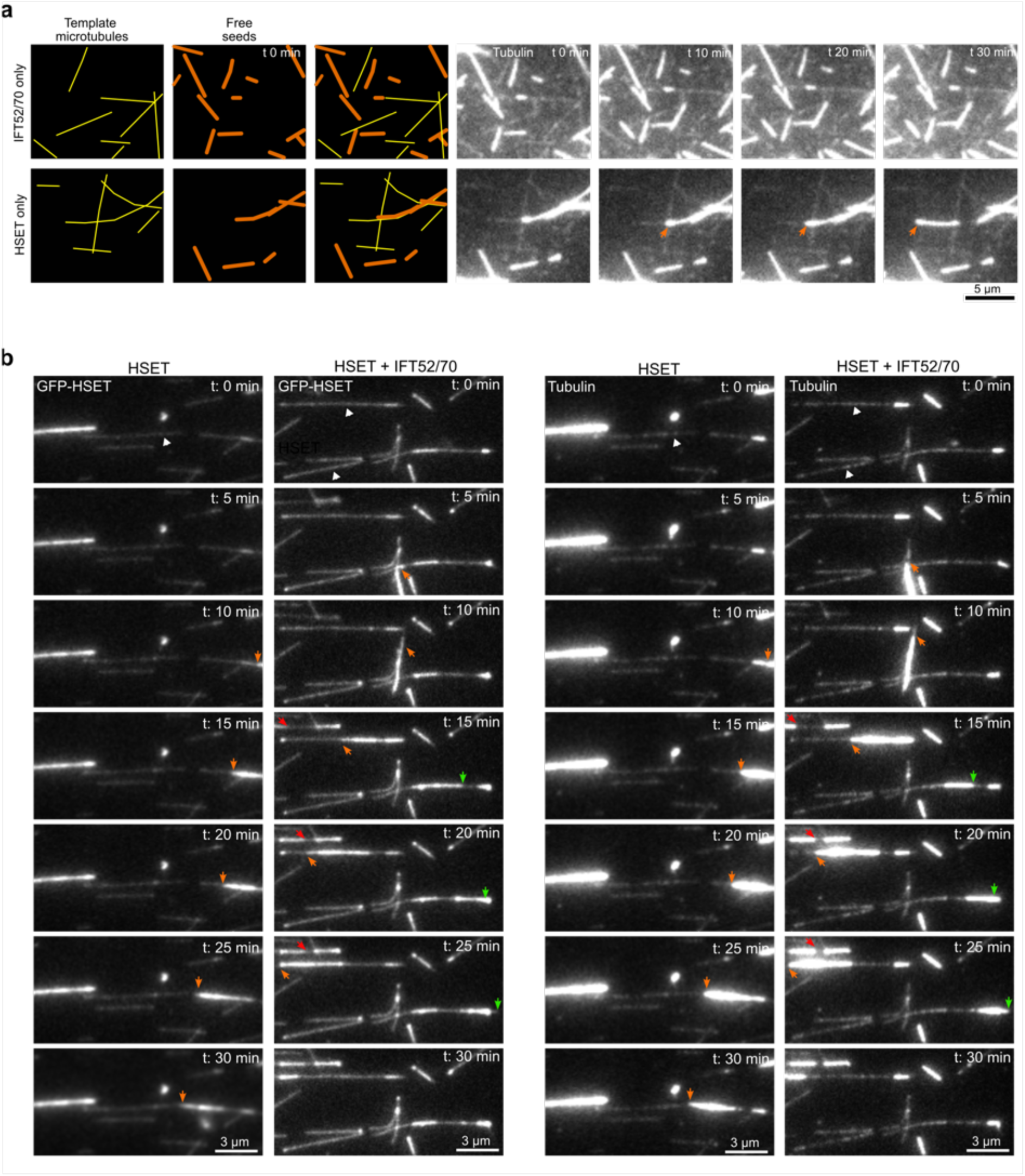
**a.** Top, schematic and representative TIRF images of free microtubule seeds and template microtubules in presence of IFT52/70 alone. No events of seeds bundling and sliding on template microtubules were observed. Bottom, schematic and representative TIRF images of free microtubules seeds bundling and sliding along template microtubules in presence of GFP-HSET. Orange arrowhead indicate a sliding seed. **b.** Still images corresponding to movies 4 and 5. The two left columns, show the GFP-HSET channel (movie 4) and the two right columns show the tubulin channel (movie 5). The free microtubule seeds are indicated with orange, red and green arrowheads. Examples of microtubule templates are indicated with white arrowheads. The conditions are: HSET alone (50 nM) or HSET (50 nM) + IFT52/70 (100 nM, right).

**Supplementary Figure 3.**
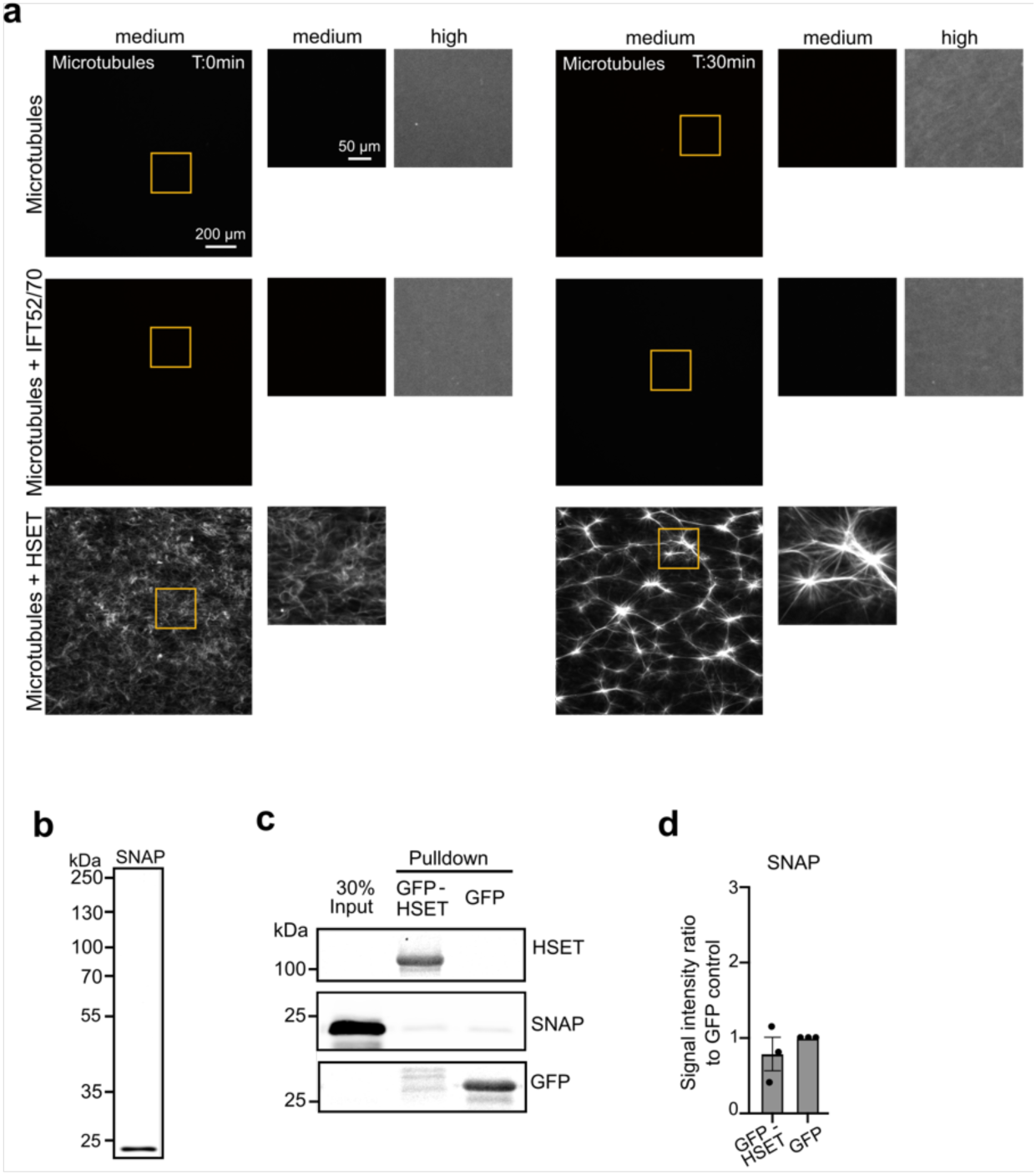
**a.** Still images from movie showing that IFT52/70 alone cannot form active microtubule networks. The positive control, HSET, displays the same images as the one shown in Fig. 5 a. Insets are the 3x magnification of the orange square in the main field of view. Medium and High refer to contrast and intensity. Even with high contrast and intensity no bundle or active network are observed with IFT52/70 alone. **b.** Coomassie blue staining of purified SNAP-tag. **c.** Western-blot of a GFP-Trap pull down of SNAP-tag. **d**. Quantification of the ratio SNAP-tag pulled-down by GFP-HSET compared to a GFP alone control. Bars represent the average of three independent experiments error bars s.e.m.

## Movies legends

**Movie 1. TIRF microscopy movie of GFP-HSET particle on microtubules, HSET + IFT52/70 condition.**

The field of view corresponds to Supplementary Fig. 1a. Time interval is 1.4 sec. The movie runs at 10 fps. The movie shows the tubulin channel (555 nm), the GFP-HSET channel (488 nm) and the IFT52/70 (647 nm) channel from top to bottom. Scale bar 3 µm.

**Movie 2. TIRF microscopy movie of GFP-HSET particle on microtubules, HSET alone condition.**

The field of view corresponds to Supplementary Fig. 1a. Time interval is 1.4 sec. The movie runs at 10 fps. The movie shows the tubulin channel (555 nm), the GFP-HSET channel (488 nm) and the 647 nm channel from top to bottom. Scale bar 3 µm.

**Movie 3. TIRF microscopy movie of GFP-HSET particle on microtubules, HSET + IFT46 condition.**

The field of view corresponds to Supplementary Fig. 1a. Time interval is 1.4 sec. The movie runs at 10 fps. The movie shows the tubulin channel (555 nm), the GFP-HSET channel (488 nm) and the IFT46 (647 nm) channel from top to bottom. Scale bar 3 µm.

**Movie 4. TIRF microscopy movie of a microtubule seed sliding on a microtubule template in absence or presence of GFP-HSET**

The field of view corresponds to Supplementary Fig. 2b left two columns. Left panel HSET alone condition, right panel HSET +IFT52/70 condition. GFP-HSET (488 nm) is visualized on this movie. Time interval is 1 min. The movie runs at 9 fps

**Movie 5. TIRF microscopy movie of a microtubule seed sliding on a microtubule template in absence or presence of GFP-HSET**

The field of view corresponds to Supplementary Fig. 2b right two columns. Left panel HSET alone condition, right panel HSET +IFT52/70 condition. Tubulin (555 nm) is visualized on this movie. Time interval is 1 min. The movie runs at 9 fps.

**Movie 6. Wide field epifluorescence microscopy movie of an active microtubule network organizing over time upon the activity of GFP-HSET.**

The field of view corresponds to Fig. 5a top row. Tubulin is visualized on this movie. Time interval is 1.5 min. The movie runs at 10 fps. Scale bar 200 µm.

**Movie 7. Wide field epifluorescence microscopy movie of an active microtubule network organizing over time upon the activity of GFP-HSET with IFT52/70.**

The field of view corresponds to Fig. 5a bottom row. Tubulin is visualized on this movie. Time interval is 1.5 min. The movie runs at 10 fps. Scale bar 200 µm.

## Notes

### Competing Interest Statement

The authors have declared no competing interest.

### Summary of Updates

Figure 1, Figure 3 and Supplementary Figure 2 are revised/updates as well as all videos

